# Threatened and Priority listed Melaleuca species from Western Australia display high susceptibility to *Austropuccinia psidii* in controlled inoculations

**DOI:** 10.1101/2023.09.19.558530

**Authors:** A. M. Martino, R. F. Park, P. A. Tobias

## Abstract

*Austropuccinia psidii* causes rust disease on species within the family Myrtaceae and was first detected in Australia in 2010, with the first detection in Western Australia in 2022. While species within the genus *Melaleuca* from Eastern Australia show variable responses to the pathogen, little is known of the response of species from Western Australia. This study established that 13 previously unscreened species of *Melaleuca*, including Threatened and Priority species that were grown from seeds sourced from Western Australian populations, were susceptible to the pandemic strain of the pathogen. The proportion of highly susceptible plants within a single species ranged from 2% – 94%, with several species displaying highly variable levels of resistance to *A. psidii*. These results highlight the importance of disease screening and may direct conservation efforts.

## Introduction

*Austropuccinia psidii*, formerly *Puccinia psidii* (Beenken 2017), is a rust fungus and the causal agent of the disease myrtle rust which impacts species within the family Myrtaceae. Originating in Brazil, the first detection of the pathogen in Australia was in 2010 and it has since spread to all states and territories except South Australia (Carnegie et al. 2010; Carnegie and Lidbetter 2012; Westaway 2016; Department of Natural Resources and Environment Tasmania 2020; Agriculture Victoria 2022; The Department of Primary Industries and Regional Development 2022a). The most recent detection within Australia was in the Kimberley region of Western Australia (WA), where infection was observed on two *Melaleuca* species near the Northern Territory border (The Department of Primary Industries and Regional Development 2022b).

*Austropuccinia psidii* infects the young and expanding tissues of susceptible hosts, including the leaves, stems, petioles, and reproductive and seed-bearing structures. In susceptible species, yellow urediniospores appear on the infected surfaces, which may be followed by other symptoms such as leaf distortion and defoliation (Pegg et al. 2014). In species with no resistance to *A. psidii*, repeated infections may lead to tree death as a result of defoliation, and impact reproduction through infection of reproductive and seed-bearing structures (Carnegie et al. 2016). In Australia, *A. psidii* has caused the near extinction of several rainforest understory species including *Rhodamnia rubescens* and *Rhodomyrtus psidioides* (Pegg et al. 2014; Carnegie et al. 2016; Environment Protection and Biodiversity Conservation Act, 1999), and could be potentially devastating for other keystone species including *Melaleuca quinquenervia* (Pegg et al. 2018).

*Melaleuca* is the third largest genus within the family Myrtaceae, comprising over 200 species (Ryan 2016) that are adapted to a range of habitats (Naidu et al. 2000). Although well adapted, changing conditions as a result of climate change are contributing to the decline of *Melaleuca* species in Australia (Saintilan et al. 2019). An increased threat is placed on these species by *A. psidii*, with several *Melaleuca* species found to be highly susceptible to the pathogen under field conditions and in controlled inoculations (Carnegie and Lidbetter 2012; Morin et al. 2012; Pegg et al. 2014, 2018; Berthon et al. 2019; Martino et al. 2022). Further, climatic modelling predicts changes in climatic suitability for the pathogen as a consequence of climate change, with increased suitability in areas of NSW, TAS, VIC, and WA (Berthon et al. 2018).

WA is rich in *Melaleuca* species, with the greatest diversity and highest level of endemism located within the South-West region of the state, with up to 72 *Melaleuca* species per 100km^2^ and endemism scores of up to 9.9 (Brophy et al. 2013). Many of these species are valued for their important ecological, cultural, and economic roles (Brophy et al. 2013). In the absence of the pathogen in the many parts of WA, the vulnerability of many *Melaleuca* species remains unknown. With the arrival of *A. psidii* into WA and high susceptibility of several *Melaleuca* species, there is an urgent need to expand current disease screening of WA species to aid pre-emptive conservation and monitoring efforts. Here, we investigated the response of 13 previously untested *Melaleuca* species to controlled inoculation with *A. psidii*. Using seed sourced from populations in areas climatically suited to *A. psidii* (Berthon et al. 2018), we aimed to determine the risk the pathogen may pose in the natural environment.

## Materials and methods

### Species selection

To determine the response of selected *Melaleuca* species from WA to *A. psidii*, seed was obtained from the Department of Biodiversity, Conservation and Attractions (DBCA) Kings Park and Kensington seed banks. Seed was obtained for species listed under the Biodiversity Conservation (BC) Act (2016) as Threatened, including critically endangered, endangered, or vulnerable species (*Biodiversity Conservation Act* 2016 (WA) s 19), and species listed as Priority on DBCA’s priority flora list (Department of Biodiversity, Conservation and Attractions 2017). While not designated under the BC Act, Priority listed species may be threatened but lack sufficient survey data to list under the Act. Seed from Priority listed species for this work include; *Melaleuca dempta, Melaleuca incana* ssp. *gingilup, Melaleuca penicula, Melaleuca similis*, and *Melaleuca sophisma*. Seed was also obtained for the Threatened (endangered) listed species *Melaleuca* sp. *Wanneroo*. Seed was also obtained from species listed as Not Threatened (*Biodiversity Conservation Act* 2016 (WA) s 19) and included *Melaleuca acutifolia, Melaleuca argentea, Melaleuca cajuputi* ssp. *cajuputi, Melaleuca fulgens* ssp. *fulgens, Melaleuca lanceolata, Melaleuca lateralis*, and *Melaleuca viminea* ssp. *appressa*. For each species, seed was collected from multiple trees with co-ordinates obtained and mapped (Figure 1).

**Figure 1.**
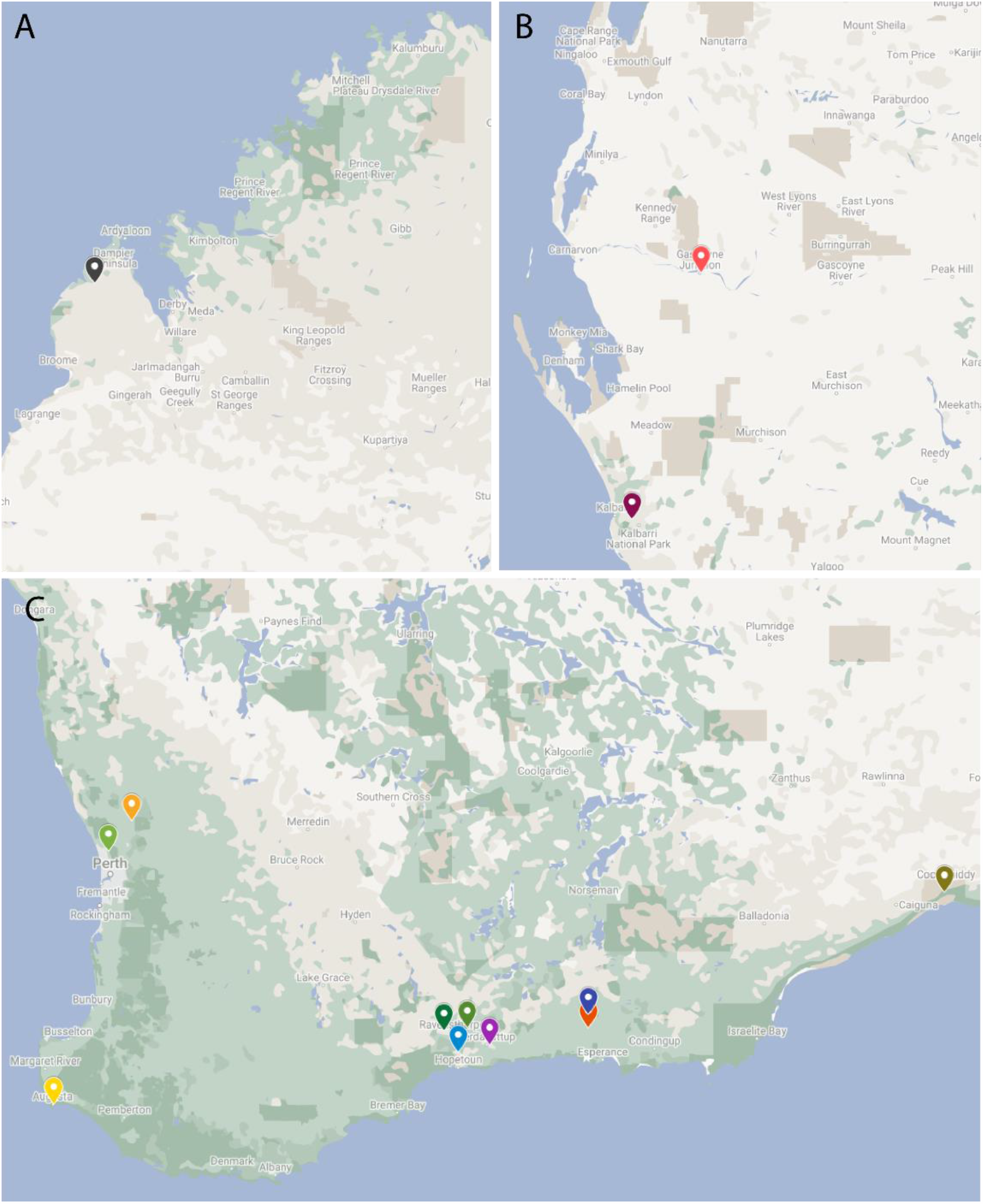
Seed collection sites by coordinate or nearest town for (A) *Melaleuca cajuputi* ssp. *cajuputi*, (B) *Melaleuca argentea* (red), *Melaleuca acutifolia* (plum), (C) *Melaleuca fulgens* ssp. *fulgens* (mustard), *Melaleuca* sp. *Wanneroo* (light green), *Melaleuca incana* ssp. *gingilup* (yellow), *Melaleuca penicula* (dark green), *Melaleuca sophisma* (light blue), *Melaleuca lateralis* (green), *Melaleuca similis* (purple), *Melaleuca viminea* ssp. *appressa* (dark blue), *Melaleuca dempta* (orange), and *Melaleuca lanceolata* (olive). Seed was collected from multiple parents at each site. Image generated in Google My Maps and interactive map is viewable at https://tinyurl.com/zvffxccd.

### Seed germination and plant growth

Seeds were sown into perforated trays containing a mix of 2:1:1 peat, coconut coir, and perlite supplemented with Osmocote® Native Controlled Release Fertiliser then covered with a fine coating of vermiculite. Perforated trays were placed into solid trays filled with 1 cm of water, every 3-4 days allowing for periods of drying to promote root growth. Seeds were germinated under natural light in a climate-controlled greenhouse set at 24°C/20°C day-time/night-time temperature on a 12 hour cycle. Germinated seedlings were transplanted into 85 mL pots (5 cm diameter and depth) containing a mix of 2:1:1 Osmocote® Native Premium Potting Mix, peat, and perlite supplemented with Osmocote® Native Controlled Release Fertiliser then placed on capillary mats. Seedlings were grown under the same light and temperature conditions as for germination.

### Seedling inoculations

For all species, we inoculated seedlings approximately 4 months post germination at the Plant Breeding Institute at the University of Sydney (Cobbitty, NSW) alongside four highly susceptible *Syzygium jambos* plants as positive controls. Approximately 50 mg of *A. psidii* urediniospores from a greenhouse increased single pustule isolate (accession 622) (Sandhu and Park 2013) was added to 50 mL of Isopar® for a final concentration of 1 mg spores/mL. Seedlings were inoculated with the suspension using an aerosol sprayer and relocated to a humid incubation chamber for 24 hours at 20°C. After incubation, seedlings were transferred to a greenhouse with the temperature set to 24°C/20°C day-time/night-time temperature on a 12-hour cycle under natural light.

### Disease susceptibility scoring

Host response to *A. psidii* inoculation was scored using a 1 – 5 scoring system based on Morin et al. (2012) and adapted for disease scoring on *Melaleuca* species (this study) where 1 indicates completely resistant or no visible response and 5 indicates highly susceptible (Table 1). *Syzygium jambos* was scored as score 5 for each round of inoculation indicating successful inoculation. As inoculations were carried out in winter under shorter day-length conditions, disease symptoms were slower to develop than in previous screening (Martino et al. 2022). Plants were left for 16 days prior to scoring to allow for complete development of plant disease symptoms.

**Table 1.**
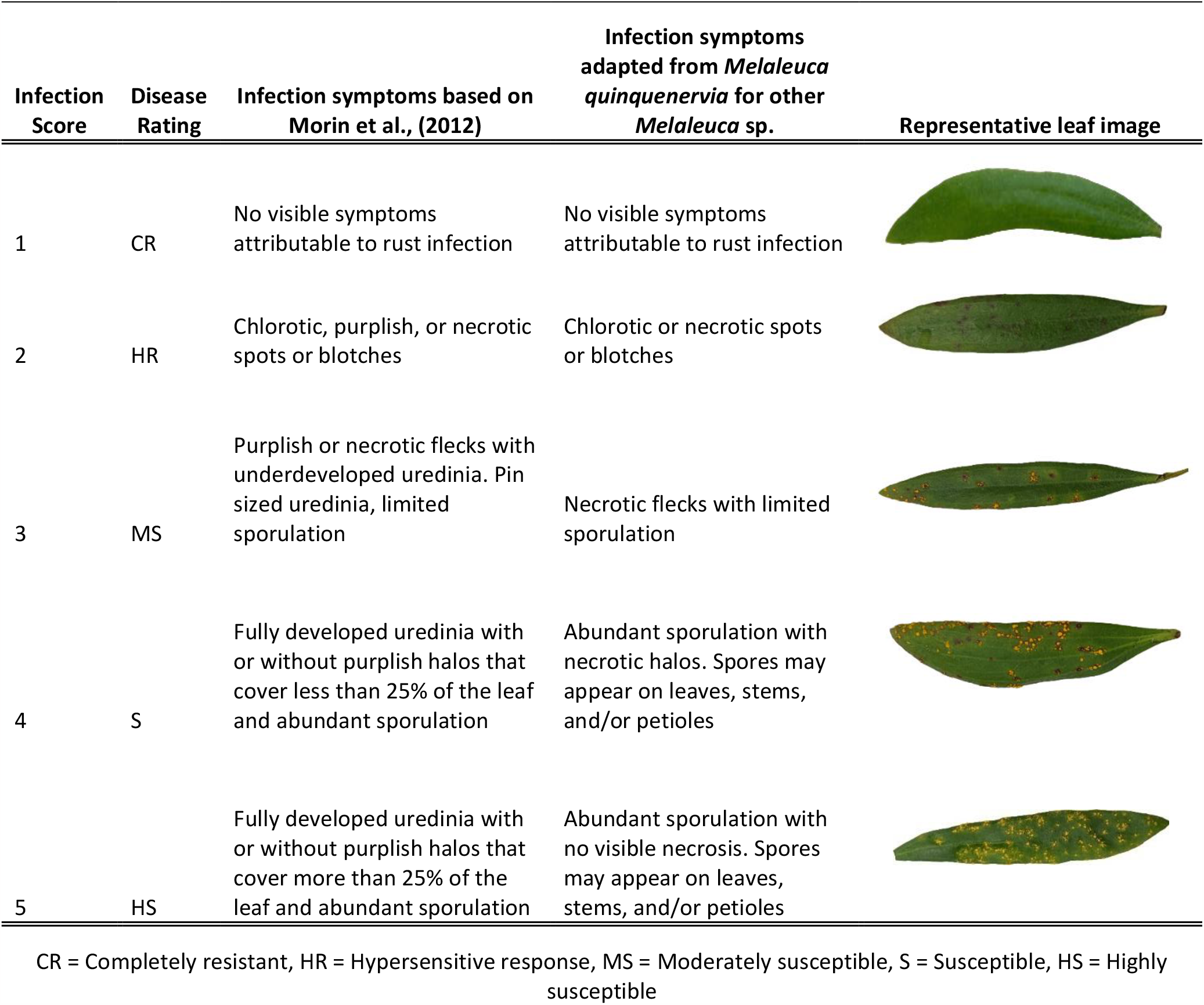
Disease scoring scale adapted from Morin et al. (2012) to score *Melaleuca* species for their response to *Austropuccinia psidii* in controlled inoculations. Scoring was based on the disease symptoms on *Melaleuca quinquenervia* scored at 14-days post inoculation with greenhouse increased single pustule isolate (accession 622) *Austropuccinia psidii* urediniospores (Sandhu and Park 2013)

| Infection Score | Disease Rating | Infection symptoms based on Morin et al., (2012) | Infection symptoms adapted from *Melaleuca quinquenervia* for other *Melaleuca* sp. | Representative leaf image |
| --- | --- | --- | --- | --- |
| 1 | CR | No visible symptoms attributable to rust infection | No visible symptoms attributable to rust infection |  |
| 2 | HR | Chlorotic, purplish, or necrotic spots or blotches | Chlorotic or necrotic spots or blotches |  |
| 3 | MS | Purplish or necrotic flecks with underdeveloped uredinia. Pin sized uredinia, limited sporulation | Necrotic flecks with limited sporulation |  |
| 4 | S | Fully developed uredinia with or without purplish halos that cover less than 25% of the leaf and abundant sporulation | Abundant sporulation with necrotic halos. Spores may appear on leaves, stems, and/or petioles |  |
| 5 | HS | Fully developed uredinia with or without purplish halos that cover more than 25% of the leaf and abundant sporulation | Abundant sporulation with no visible necrosis. Spores may appear on leaves, stems, and/or petioles |  |

CR = Completely resistant, HR = Hypersensitive response, MS = Moderately susceptible, S = Susceptible, HS = Highly susceptible

### Imaging

All images were captured using an Olympus OM-5 fitted with an Olympus M. Zuiko Premium 60mm f/2.8 Macro Lens. Raw images were processed using Adobe Photoshop 2023.

## Results

Within 16 days post inoculation, symptoms had developed on the highly susceptible *S. jambos* control plants (Figure 2 A-B). Using the scoring system adapted for *Melaleuca* species (this study, Table 1), disease scores across all plants ranged from completely resistant (score 1) to highly susceptible (score 5) (Supplementary Figures 1 – 13). The proportion of highly susceptible plants within a single species ranged from 2% for *M. sophisma*, to 94% for *M. lateralis*. Eight of the species had individuals that were either completely resistant (score 1) or highly susceptible (score 4 or 5) to *A. psidii* (Table 2). This binary response was observed for *M. lateralis*, with 94% of plants highly susceptible and 6% with no observable symptoms and for *M. sophisma*, with 2% of plants highly susceptible and the remaining 98% with no observable symptoms (Table 2). Only three of the 13 species tested; *M. argentea, M. cajupti* spp. *cajupti*, and *M. incana* ssp. *gingilup*, had representative plants from each disease score (Table 2). Urediniospores were observed on the leaves of all highly susceptible plants (Figure 2 C-O), as well as infection on stems and petioles on all species except for *M. sophisma* and *M. incana* ssp. *gingilup*.

**Table 2.** Disease scoring, based on Morin et al. (2012) and adapted for *Melaleuca* species (this study), of controlled inoculation of *Austropuccinia psidii* of Threatened (*Biodiversity Conservation Act* 2016 (WA) s 19) and Priority (Department of Biodiversity, Conservation and Attractions 2017) listed *Melaleuca* species included total number of plants scored and the percentage of plants observed in each disease scoring category

| Species name | Listing under Biodiversity Conservation Act 2016 or DBCA Priority Flora List | Total Number of Plants Scored | Disease Score (% of Total Plants) |  |  |  |  |
| --- | --- | --- | --- | --- | --- | --- | --- |
|  |  |  | 1 | 2 | 3 | 4 | 5 |
| *Melaleuca acutifolia* | NT | 20 | 10 | 0 | 0 | 20 | 70 |
| *Melaleuca argentea* | NT | 52 | 44 | 23 | 4 | 4 | 25 |
| *Melaleuca cajuputi* ssp. *cajuputi* | NT | 54 | 28 | 15 | 13 | 35 | 9 |
| *Melaleuca dempta* | P3 | 11 | 9 | 0 | 0 | 0 | 91 |
| *Melaleuca fulgens* ssp. *fulgens* | NT | 27 | 18 | 0 | 18 | 7 | 57 |
| *Melaleuca incana* ssp. *gingilup* | P2 | 41 | 32 | 5 | 5 | 19 | 39 |
| *Melaleuca lanceolata* | NT | 27 | 15 | 0 | 18 | 26 | 41 |
| *Melaleuca lateralis* | NT | 31 | 6 | 0 | 0 | 0 | 94 |
| *Melaleuca penicula* | P4 | 32 | 66 | 0 | 0 | 6 | 28 |
| *Melaleuca similis* | P1 | 45 | 24 | 0 | 0 | 0 | 76 |
| *Melaleuca sophisma* | P1 | 55 | 98 | 0 | 0 | 0 | 2 |
| *Melaleuca* sp. *Wanneroo* | EN | 80 | 14 | 0 | 0 | 0 | 86 |
| *Melaleuca viminea* ssp. *appressa* | P2 | 25 | 28 | 0 | 0 | 4 | 68 |

EN = Endangered (Threatened species considered to be “facing a very high risk of extinction in the wild in the near future, as determined in accordance with criteria set out in the ministerial guidelines”), NT = Not Threatened, P1 - 3 = Priority 1 – 3 (Poorly-known species with conservation threat highest for Priority 1), P4 = Priority 4 (Rare, Near Threatened and other species in need of monitoring).

**Figure 2.**
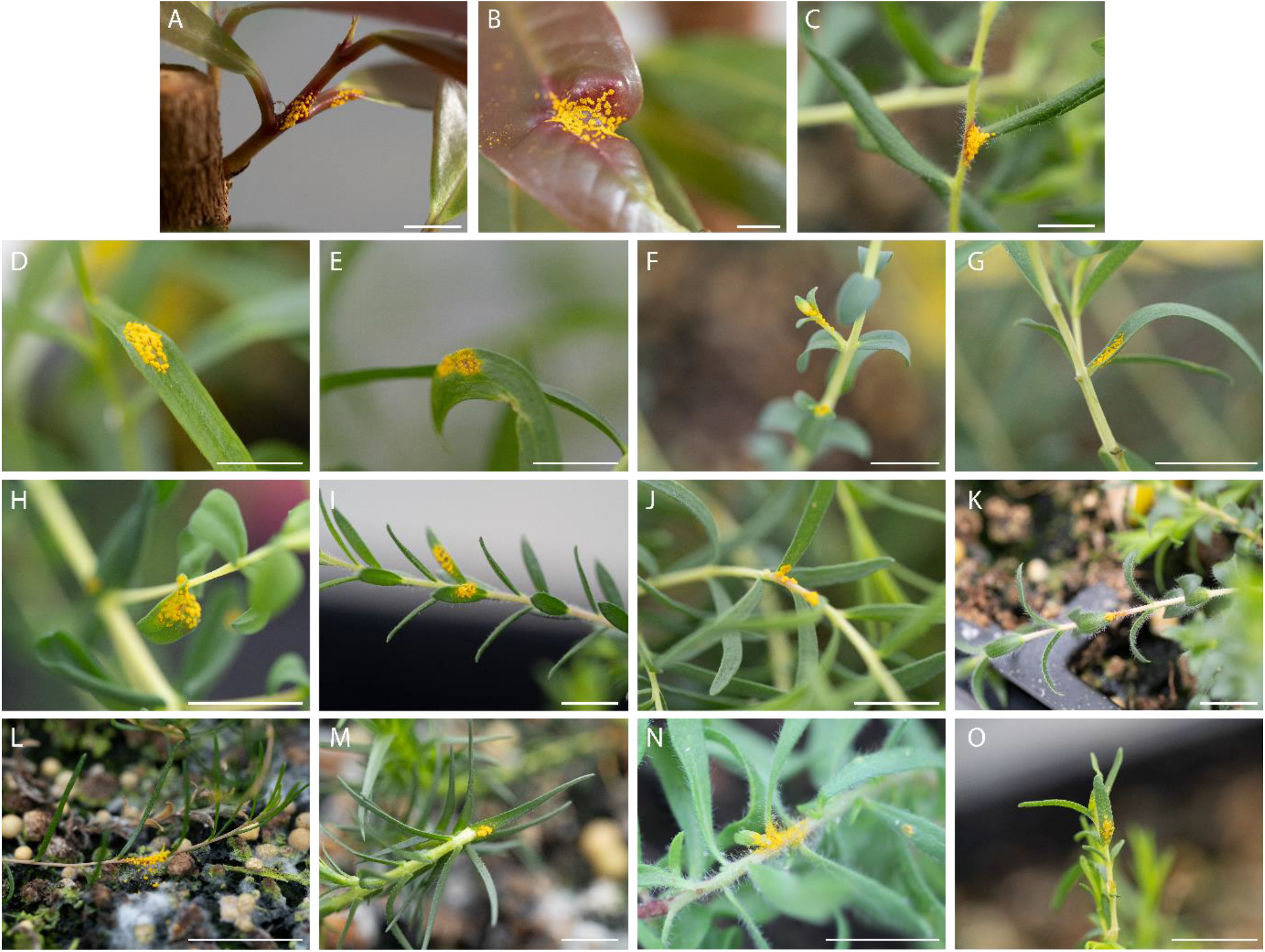
Representative highly susceptible disease symptoms on (A – B) *Syzygium jambos* positive control, (C) *Melaleuca acutifolia*, (D) *Melaleuca argentea*, (E) *Melaleuca cajuputi* ssp. *cajuputi*, (F) *Melaleuca dempta*, (G) *Melaleuca fulgens* ssp. *fulgens*, (H) *Melaleuca incana* ssp. *gingilup*, (I) *Melaleuca lanceolata*, (J) *Melaleuca lateralis*, (K) *Melaleuca penicula*, (L) *Melaleuca similis*, (M) *Melaleuca sophisma*, (N) *Melaleuca* sp. *Wanneroo*, and (O) *Melaleuca viminea* ssp. *appressa*. Scale bar = 0.5 cm.

## Discussion

Here, we investigated host response to *A. psidii* in 13 previously unscreened *Melaleuca* species from a range of geographic locations in Western Australia, revealing varying proportions of highly susceptible plants within and between species. The broad-leaved paperbark species included in this study, *M. cajuputi* ssp. *cajuputi* and *M. argentea*, both displayed variability in response to *A. psidii* with plants displaying symptoms in each disease scoring category. This has previously been shown for other broad-leaved species including *M. quinquenervia, M. viridiflora*, and *M. leucadendra* (Pegg et al. 2018; Martino et al. 2022). Pegg et al. (2018) assessed the proportion of resistant *M. viridiflora* from two provenances in WA determining 22-23 % of seedlings were resistant to *A. psidii*. The same study assessed *M. leucadendra* seedlings from three provenances in WA determining 1-53% resistant seedlings, while a separate study assessing a population from the Wunaamin Conservation Park in WA determined 30% of plants to be resistant to *A. psidii* (Martino et al. 2022). These results indicate variability in host response to the pathogen between populations of broad-leaved paperbarks. As the *M. cajuputi* ssp. *cajuputi* and *M. argentea* screened in this study were from grown from seed collected from a single provenance, further studies should be conducted to determine variation in host response between populations. Such information may be informative to shed light on the forces driving differences in disease resistance between populations and indicating that they may be useful for differential pathotype trials going forward.

Unlike the broad-leaved paperbarks, most species tested in this study displayed little variability in response to the pathogen. This difference may be explained by the geographic distribution differences of these species. For the broad-leaved paperbark species screened in this and in previous studies, populations are numerous, and broadly distributed across large geographic regions of Australia (Brophy et al. 2013). This distribution pattern is also true for *M. fulgens* ssp. *fulgens* and *M. lanceolata* (Western Australian Herbarium) which both display similar variability in response to the pathogen as the broad-leaved species. Conversely for *M. dempta, M. penicula, M. similis, M. sophisma, M*. sp. *Wanneroo*, and *M. viminea* ssp. *appressa* where populations are geographically sparse (Western Australian Herbarium), all display low variability in pathogen response. As *Melaleuca* species are predominantly outcrossing (Quang Tan 2008; Baskorowati et al. 2010; Brophy et al. 2013; Kartikawati et al. 2021), these differences may be explained by reductions in gene flow within small, isolated populations, resulting in reduced genetic diversity within populations.

Of particular interest is the high proportion of resistant *M. sophisma* plants observed, with only 2% of total plants susceptible to *A. psidii*. The remaining 98% of plants displayed no observable symptoms or hypersensitive response, potentially indicating preformed resistance mechanisms. The lack of a hypersensitive response has been observed in other Myrtaceae species inoculated with *A. psidii*, including several *Eucalyptus* species (Dos Santos et al. 2019). In species with no observable symptoms post inoculation, *A. psidii* was not detected within leaf tissues as determined by qPCR (Dos Santos et al. 2019). The results indicated that the leaves were not colonised by the pathogen, with the tested hypothesis that chemical compounds within cuticular waxes provide preformed resistance in these species (Dos Santos et al. 2019). Leaf epidermal appendages have also been implicated in contributing to responses to the pathogen with studies correlating rust susceptibility with increased trichome density (Wang et al. 2020; Varma et al. 2023). Here, the suggestion is that trichomes facilitate increased adherence of spores to the leaf surface. The lack of a hypersensitive response in 98% of unaffected *M. sophisma* plants may indicate reduced urediniospores adherence to the leaf surface owing to the absence of trichomes, or the inability of *A. psidii* to penetrate or colonise the leaves of this species owing to cuticular waxes. As many of these species remain poorly characterised, histological analyses during rust infection may shed light on preformed resistance mechanisms on these species.

Correlating disease responses in our greenhouse seedling tests with field responses will be important in defining potential risks to these species. We were encouraged by the presence of some resistant individuals within some of the listed Priority species. The results highlight the importance of continued disease screening to determine the vulnerability of individual Myrtaceae species to *A. psidii*. The identification of species with high susceptibility to the pathogen will be useful to inform disease surveillance in the natural environment and to direct conservation efforts such as seed collection.

## Acknowledgements

This work was funded by the Australian Research Council under linkage project LP190100093 and AMM by the Australian Government Research Training Program. We thank the Department of Biodiversity, Conservation and Attractions (Kings Park and Kensington) for suppling the seed for this work and Bob Makinson for the introductions that facilitated this work.

