## Supplementary Figures 1 - 13 for "Threatened and Priority listed Melaleuca species from Western Australia display high susceptibility to *Austropuccinia psidii* in controlled inoculations"

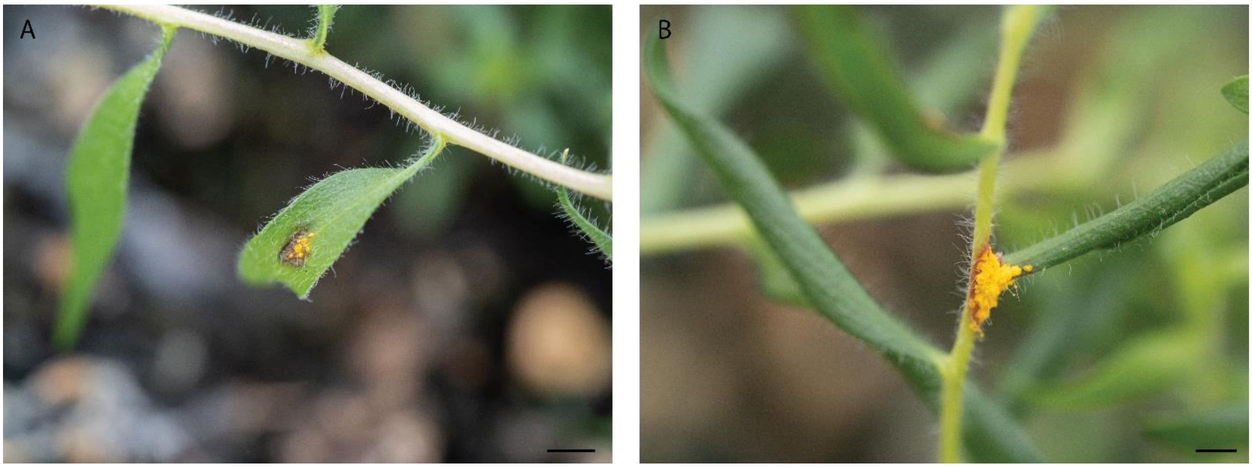


Figure 1. *Austropuccinia psidii* disease symptoms on *Melaleuca acutifolia*. Plants were assessed for disease symptoms at 16-days post inoculation and visible symptoms ranged from score 4 (A) and score 5 (B). Scale bar = 0.5cm.

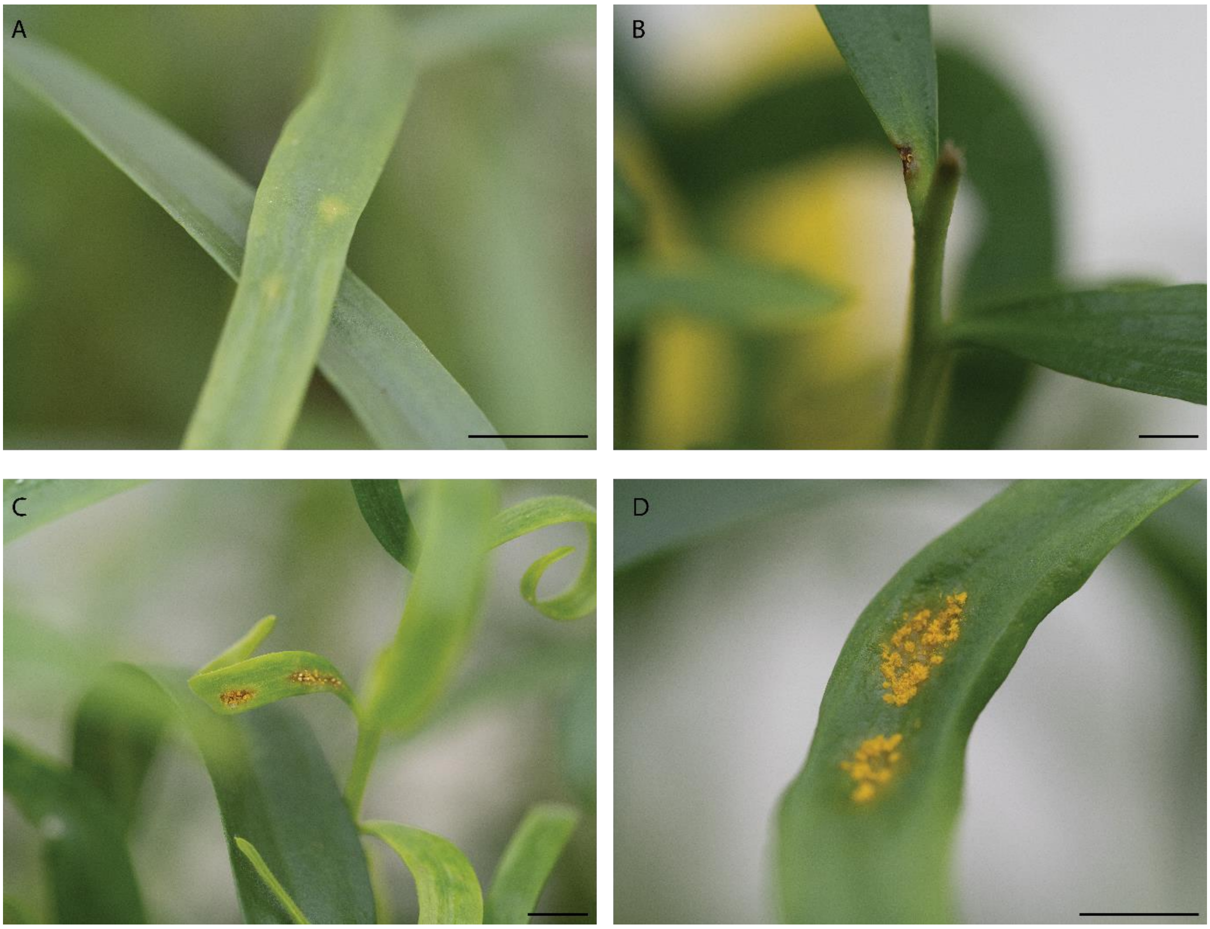


Figure 2. *Austropuccinia psidii* disease symptoms on *Melaleuca argentea*. Plants were assessed for disease symptoms at 16-days post inoculation and visible symptoms ranged from score 2 (A), score 3 (B), score 4 (C), and score 5 (D). Scale bar = 0.5cm.

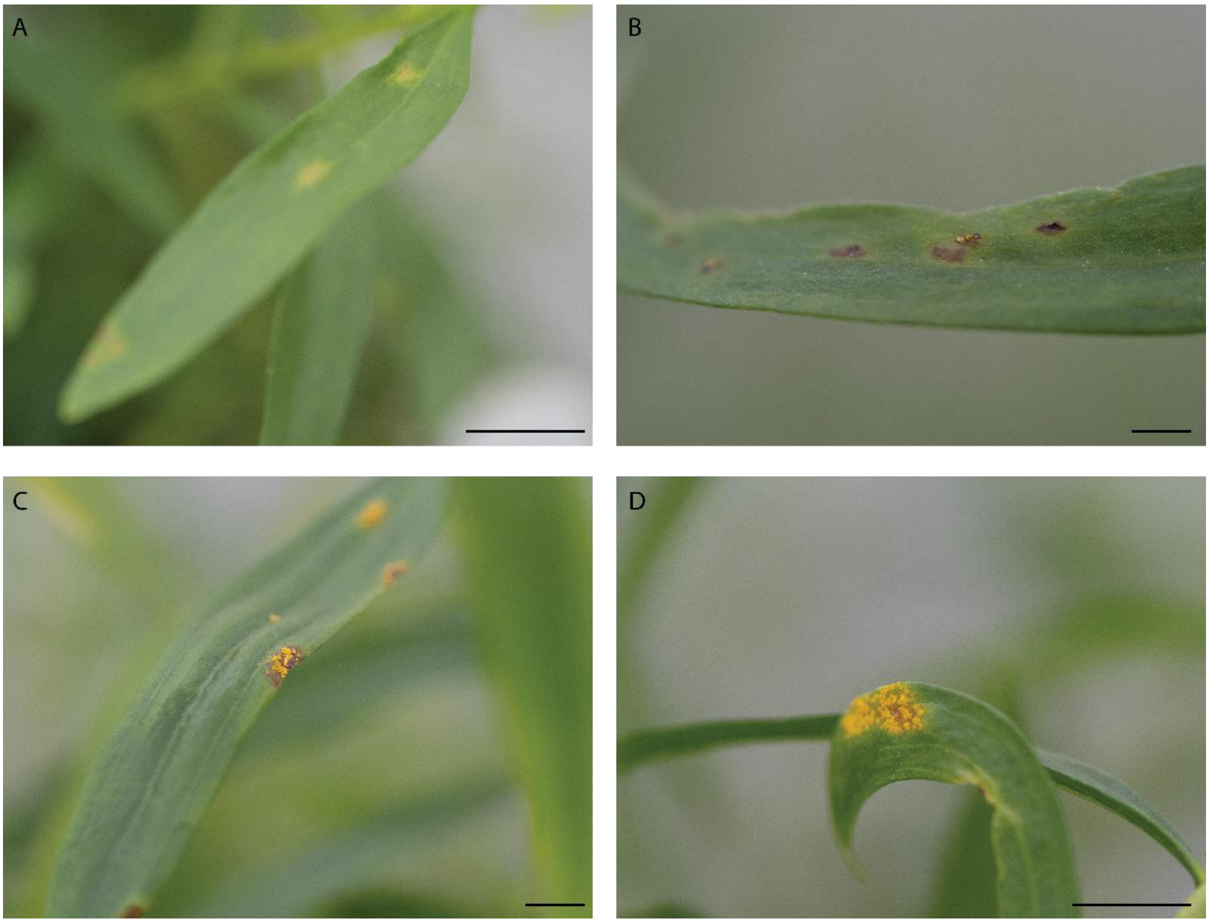


Figure 3. *Austropuccinia psidii* disease symptoms on *Melaleuca cajuputi ssp. cajuputi*. Plants were assessed for disease symptoms at 16-days post inoculation and visible symptoms ranged from score 2 (A), score 3 (B), score 4 (C), and score 5 (D). Scale bar = 0.5cm.

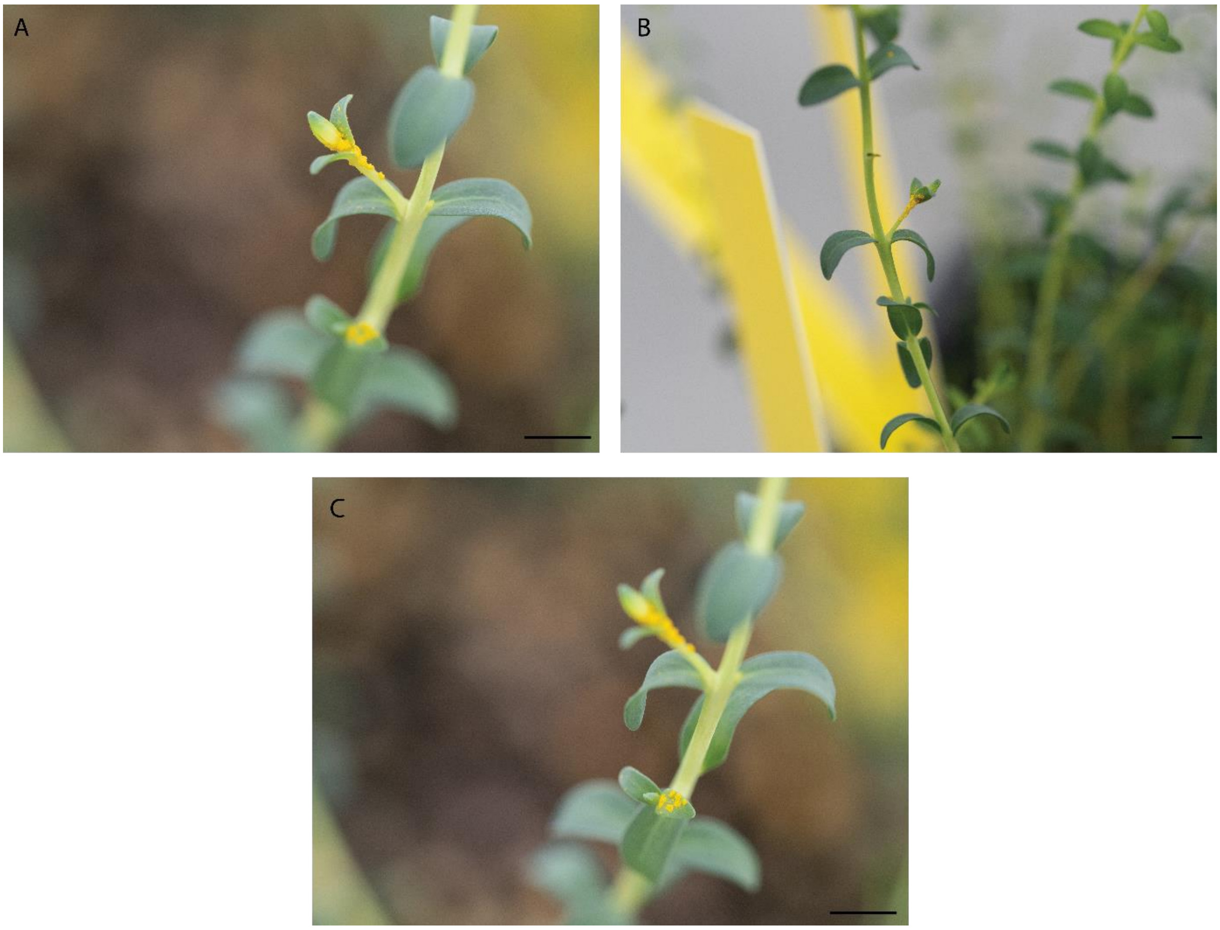


Figure 4. *Austropuccinia psidii* disease symptoms on *Melaleuca dempta*. Plants were assessed for disease symptoms at 16-days post inoculation and all plants with visible disease symptoms were scored at score 5 (A-C). Scale bar = 0.5cm.

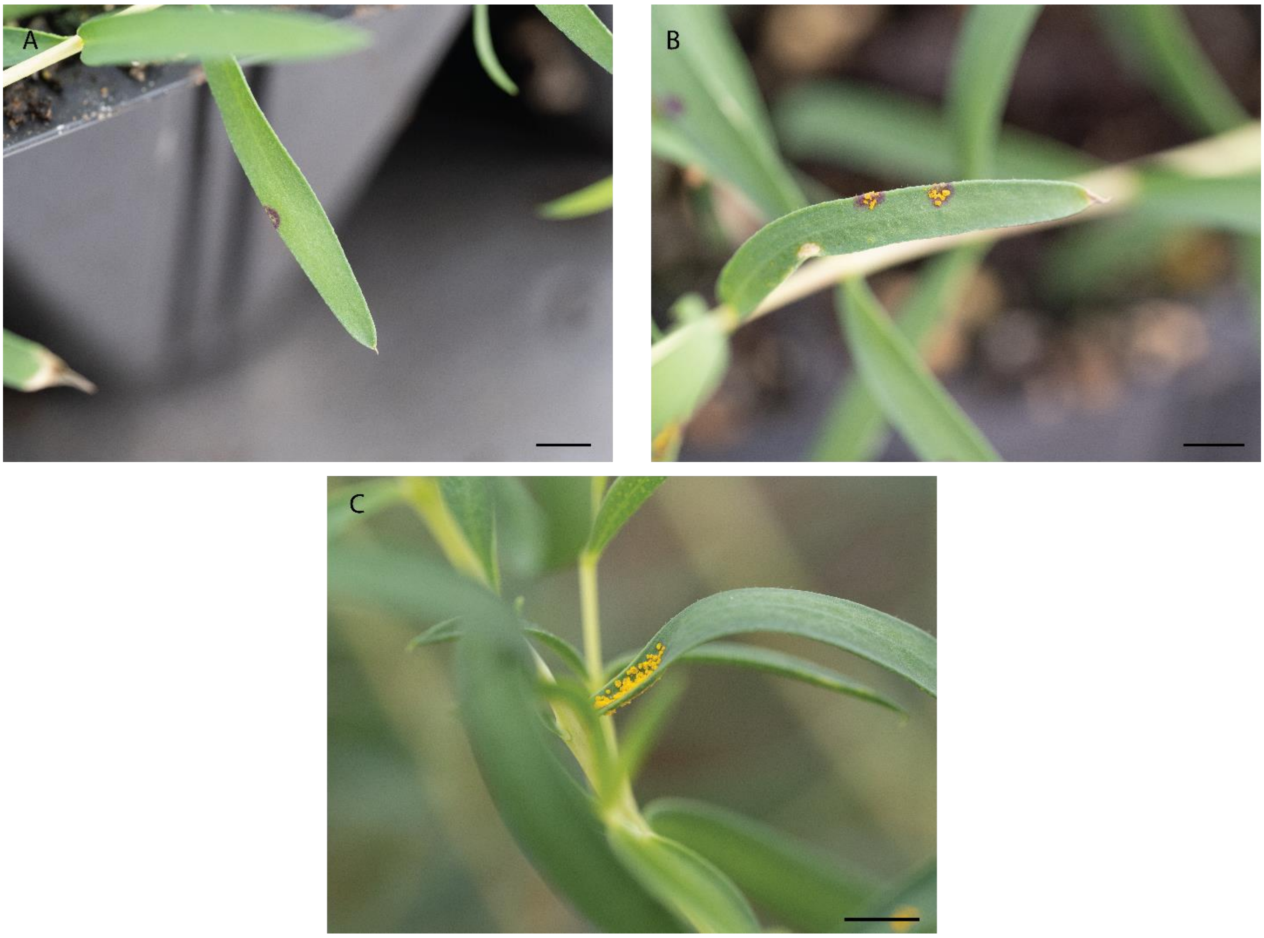


Figure 5. *Austropuccinia psidii* disease symptoms on *Melaleuca fulgens* ssp. *fulgens*. Plants were assessed for disease symptoms at 16-days post inoculation and visible symptoms ranged from score 3 (A), score 4 (B), and score 5 (C). Scale bar = 0.5cm.

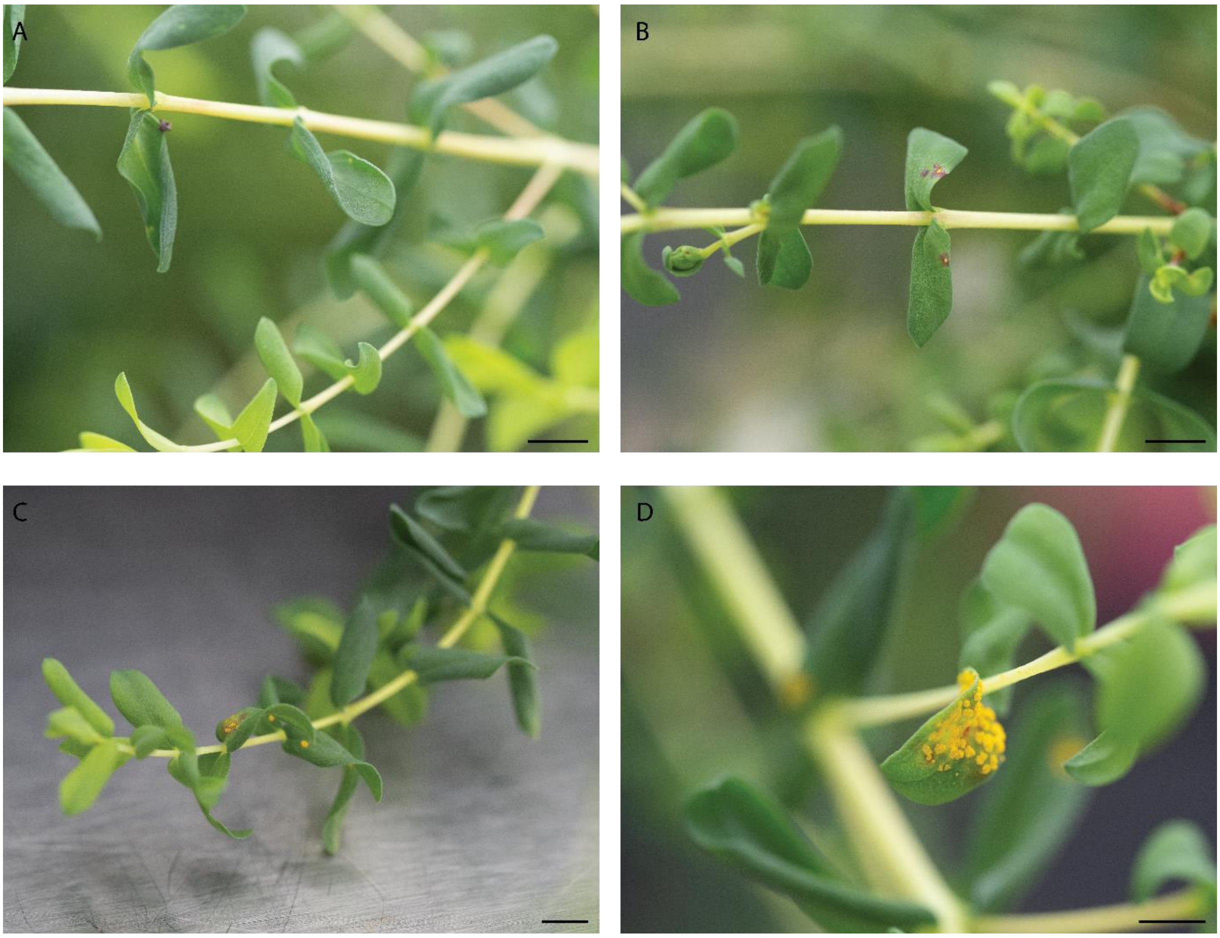


Figure 6. *Austropuccinia psidii* disease symptoms on *Melaleuca incana* ssp. *gingilup*. Plants were assessed for disease symptoms at 16-days post inoculation and visible symptoms ranged from score 2 (A), score 3 (B), score 4 (C), and score 5 (D). Scale bar = 0.5cm.

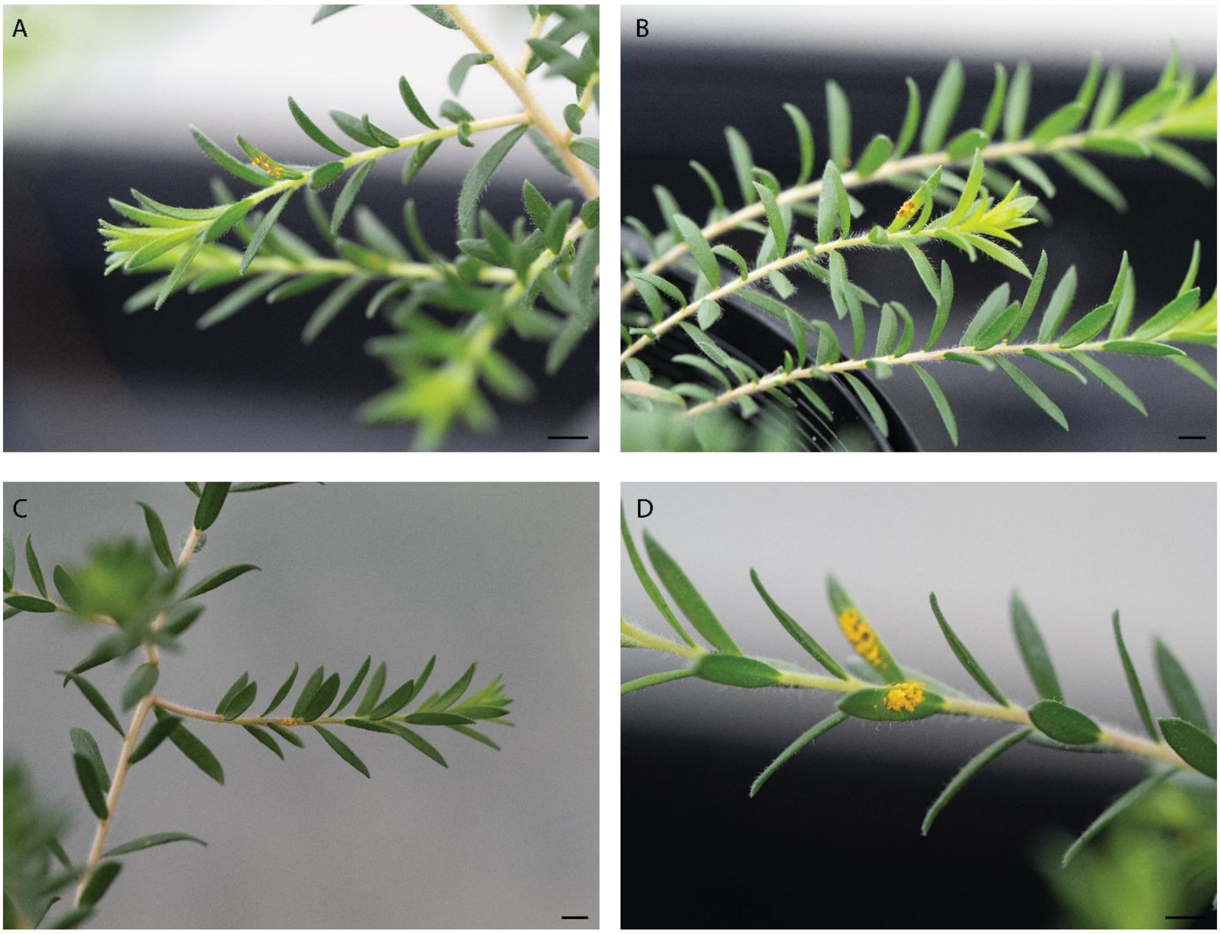


Figure 7. *Austropuccinia psidii* disease symptoms on *Melaleuca lanceolata*. Plants were assessed for disease symptoms at 16-days post inoculation and visible symptoms ranged from score 3 (A), score 4 (B), and score 5 (C & D). Scale bar = 0.5cm.

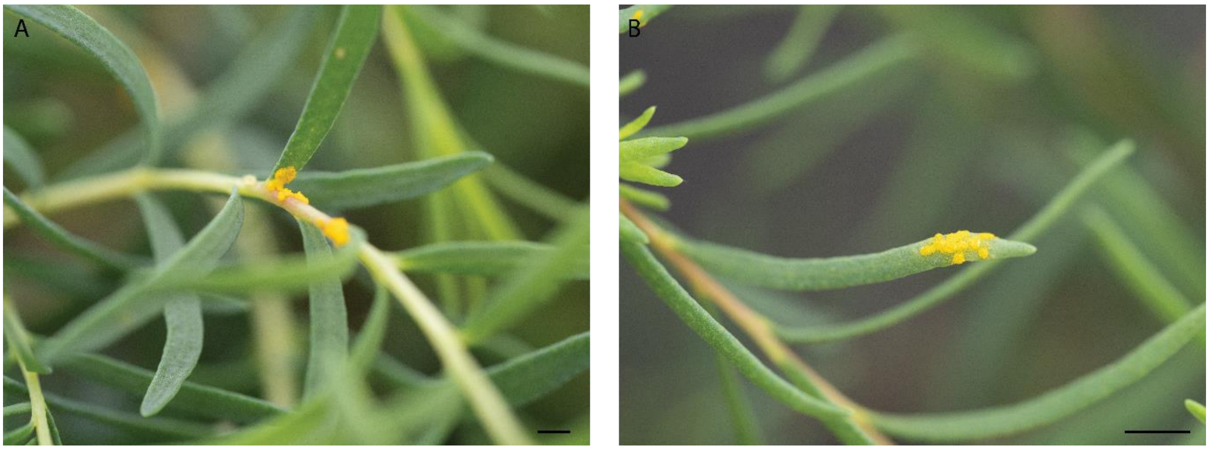


Figure 8. *Austropuccinia psidii* disease symptoms on *Melaleuca lateralis*. Plants were assessed for disease symptoms at 16-days post inoculation and all plants with visible disease symptoms were scored at score 5 (A & B). Scale bar = 0.5cm.

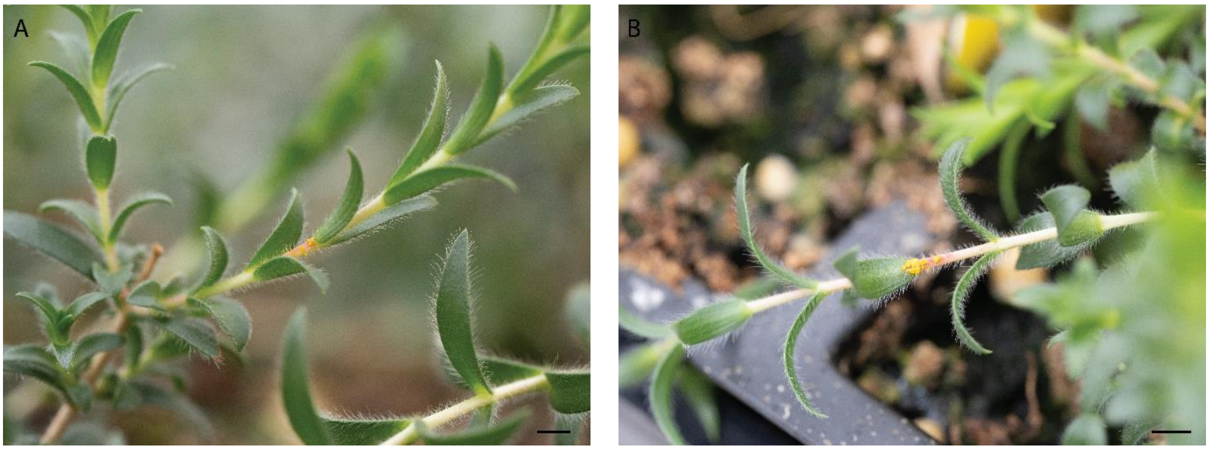


Figure 9. *Austropuccinia psidii* disease symptoms on *Melaleuca penicula*. Plants were assessed for disease symptoms at 16-days post inoculation and visible symptoms ranged from score 4(A), and score 5 (B). Scale bar = 0.5cm.

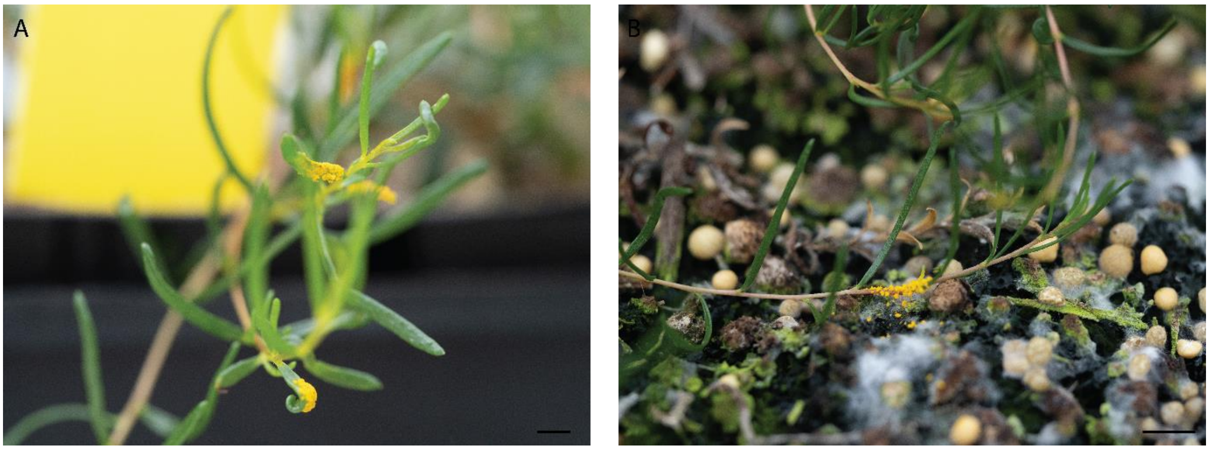


Figure 10. *Austropuccinia psidii* disease symptoms on *Melaleuca similis*. Plants were assessed for disease symptoms at 16-days post inoculation and all plants with visible disease symptoms were scored at score 5 (A & B). Scale bar = 0.5cm.

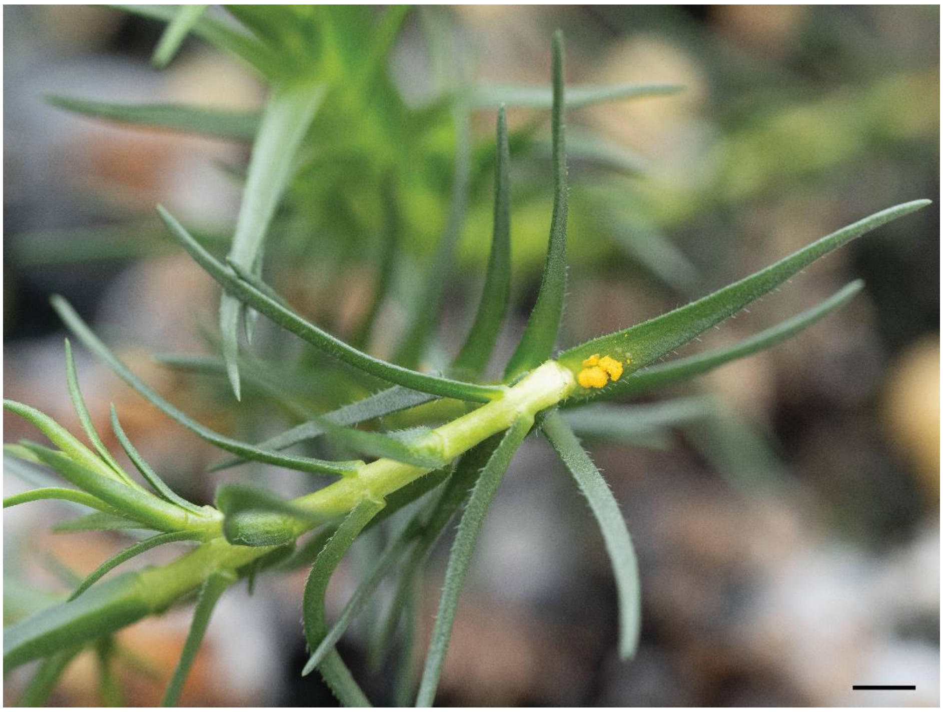

Figure 11. *Austropuccinia psidii* disease symptoms on *Melaleuca sophisma*. Plants were assessed for disease symptoms at 16-days post inoculation and all plants with visible disease symptoms were scored at score 5. Scale bar = 0.5cm.

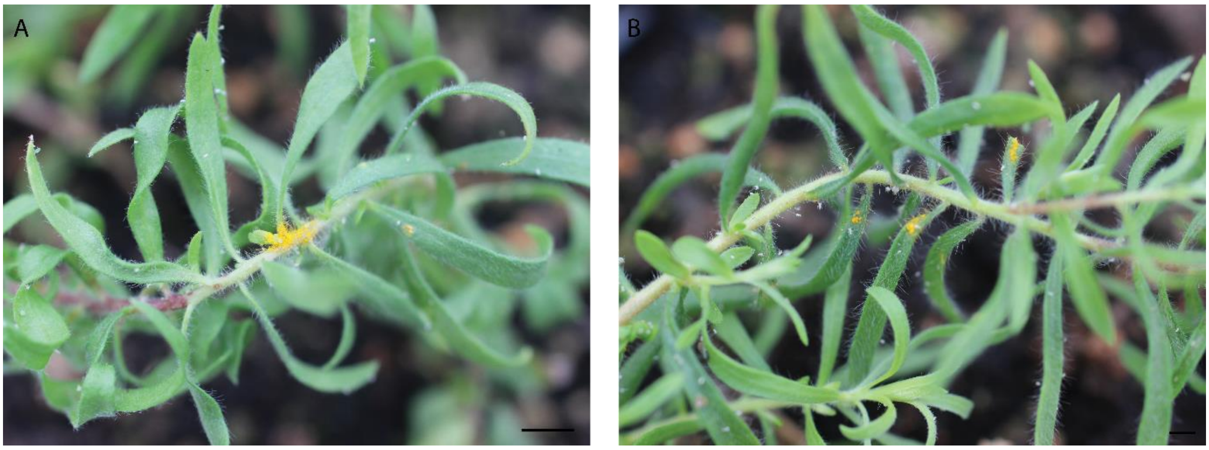


Figure 12. *Austropuccinia psidii* disease symptoms on *Melaleuca* sp. *Wanneroo*. Plants were assessed for disease symptoms at 16-days post inoculation and all plants with visible disease symptoms were scored at score 5 (A & B). Scale bar = 0.5cm.

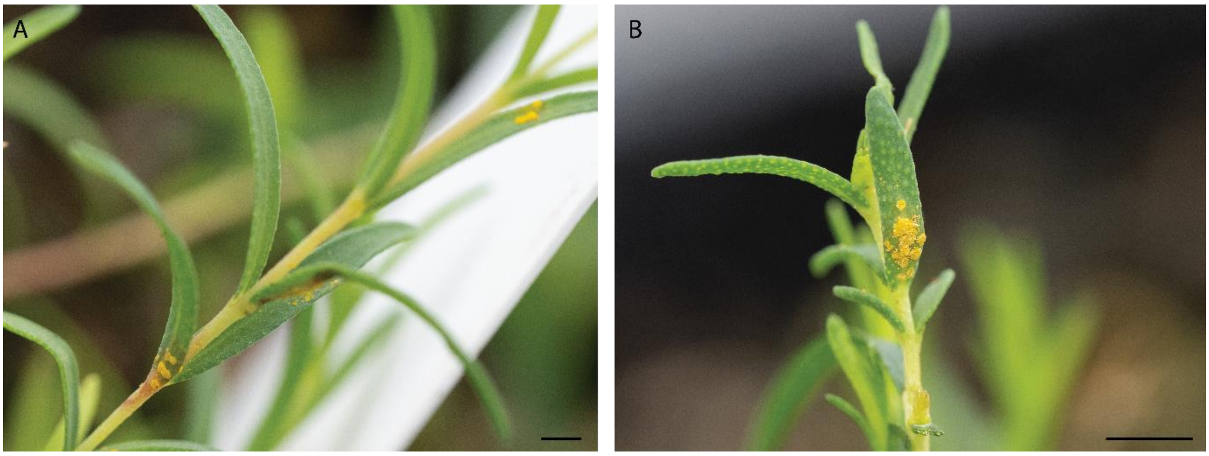


Figure 13. *Austropuccinia psidii* disease symptoms on *Melaleuca viminea* spp. *appressa*. Plants were assessed for disease symptoms at 16-days post inoculation and all plants with visible disease symptoms were scored at score 4 (A), and score 5 (B). Scale bar = 0.5cm.
